# Joint enzyme-reaction retrieval and catalytic optima prediction via multimodal fusion

**DOI:** 10.64898/2026.05.19.726405

**Authors:** Yueyi Cai, Feng Yang, Juan Liu

## Abstract

**Motivation:** Enzyme-reaction retrieval is increasingly used to prioritize candidate biocatalysts for experimental follow-up, where useful recommendations should indicate not only whether an enzyme can catalyze a target reaction but also under which pH and temperature conditions it should be tested. Existing retrieval models optimize catalytic matching scores, whereas catalytic optima predictors are typically developed as enzyme-level regressors because public pH and temperature annotations are sparse and often available only at the enzyme or EC-associated record level. This separation leaves a practical gap: high-ranking enzyme-reaction pairs are not evaluated for condition suitability, and enzyme-level optima predictions do not use the reaction context being retrieved.

**Results:** We present GERO, a multimodal fusion framework that uses feature-gated cross-modal fusion to integrate global enzyme sequence semantics, sequence-derived pocket geometry, and molecular reaction representations for condition-aware enzyme-reaction retrieval and catalytic optima estimation with reaction context. To evaluate this setting, we define the tolerance-restricted hit rate (Hit@k-TR), which requires both top-k retrieval of the correct candidate and condition prediction within predefined tolerances. Across enzyme- and reaction-similarity splits, GERO improves Hit@k-TR over two-stage retrieval-then-prediction baselines. Representative benchmark examples and an iodinin biosynthesis case study further illustrate GERO’s ability to provide candidate rankings together with plausible assay-condition estimates for downstream experimental prioritization.

**Availability and implementation:** Source code is available at https://github.com/ykxhs/GERO.

**Supplementary information:** Supplementary data are available at XXXX online.

## Introduction

Enzymes catalyze diverse biochemical transformations and are widely used in synthetic biology, pharmaceutical manufacturing and green chemistry (Kirk et al., 2002; Hemalatha et al., 2013). For computational biocatalyst discovery, a useful recommendation must answer two related questions. First, is the enzyme likely to catalyze the target reaction? Second, under what pH and temperature should the candidate be assayed? The first question concerns catalytic matching, whereas the second concerns catalytic-condition estimation. Both are important for experimental follow-up because pH and temperature strongly influence enzyme activity and stability (Fromm, 1975; Almeida and Marana, 2019).

Most existing computational methods focus on one of these questions. Enzyme function annotation and substrate-specificity models infer catalytic roles from sequence, structure or enzyme-substrate representations. For example, CLEAN uses contrastive protein representations for EC-number prediction (Yu et al., 2023), whereas EZSpecificity uses enzyme-pocket and substrate representations to model substrate specificity (Cui et al., 2025). More recently, enzyme-reaction retrieval has framed catalytic matching as a cross-modal ranking task. CLIPZyme (Mikhael et al., 2024), ReactZyme (Hua et al., 2024) and CREEP, adapted from CARE (Yang et al., 2024), rank candidate reactions for a query enzyme or candidate enzymes for a query reaction by aligning enzyme and reaction representations. These approaches address catalytic matching, but their retrieval scores do not by themselves provide pH and temperature estimates for the ranked pairs. In parallel, catalytic optima prediction has usually been formulated as an enzyme-level regression problem. EpHod predicts optimal pH from protein language-model representations (Gado et al., 2025), and Seq2Topt estimates temperature and pH optima from amino acid sequence features (Qiu et al., 2025). These models are valuable for annotating enzyme-level properties, but they are typically developed separately from enzyme-reaction retrieval and do not explicitly use the reaction being retrieved.

The observed catalytic optimum may vary with substrates, cofactors, assay protocols and reaction contexts, which can affect binding, stability and reaction energetics (Qiu et al., 2025). Practical biocatalyst discovery therefore benefits from a condition-aware formulation in which a successful recommendation should both match an enzyme to a target reaction and provide useful starting estimates of its catalytic conditions. A straightforward solution is to sequentially cascade a retrieval module with a property prediction model, forming a two-stage retrieval-then-prediction pipeline. However, this strategy does not directly optimize the practical output needed by experimental users. The retrieval module is trained to rank catalytic matches without regard to condition accuracy, whereas a separate enzyme-level predictor estimates optima without conditioning on the reaction being retrieved. The resulting objective mismatch can be compounded by cascading errors, where mistakes from the retrieval stage directly affect subsequent condition prediction (Leaman and Lu, 2016).

In this paper, we present GERO, a framework that models catalytic matching and catalytic condition prediction as a unified computational task. GERO enables bidirectional retrieval between enzymes and reactions while simultaneously estimating pH and temperature optima for retrieved candidates using the paired reaction as contextual information. For this formulation, GERO combines global evolutionary sequence information from ESM-2 (Lin et al., 2022), local geometric structures of predicted catalytic pockets derived from AlphaFold (Jumper et al., 2021) and identified by P2Rank (Krivák and Hoksza, 2018), and molecular-level reaction representations encoded from reactants and products using Uni-Mol (Zhou et al., 2023). This design reflects the complementary roles of global functional semantics and local structural context, because substrate recognition and physicochemical constraints associated with catalytic conditions can depend on the pocket environment as well as on sequence-level function. A feature-gating fusion module then constructs a joint enzyme-reaction representation for both matching-score prediction and catalytic optima estimation. To evaluate this joint task, we introduce the tolerance-restricted hit rate (Hit@k-TR), which requires the correct enzyme or reaction to appear among the top-ranked candidates and further requires predicted pH and temperature to fall within predefined tolerance ranges of the annotated optima. Benchmark evaluations show that GERO improves Hit@k-TR compared with two-stage baseline models, and an iodinin biosynthesis case study demonstrates its practical utility in prioritizing pathway-relevant enzymes while suggesting likely assay conditions for subsequent in vitro validation.

## Methods

### Task formulation

Given an enzyme-reaction pair (*e, r*), the goal is to estimate catalytic compatibility and predict catalytic pH and temperature optima conditioned on the paired reaction. GERO therefore predicts a matching score, an optimal pH estimate and an optimal temperature estimate:

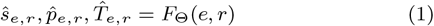

where *ŝ*_*e,r*_ is used to rank candidates, and 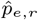 and 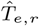 provide pH and temperature estimates for the same candidate pair. This formulation supports both enzyme-to-reaction retrieval (E→R), where a query enzyme is ranked against candidate reactions, and reaction-to-enzyme retrieval (R→E), where a query reaction is ranked against candidate enzymes. In both directions, each scored candidate pair is accompanied by pH and temperature estimates with reaction context.

### Overview of GERO

The framework combines enzyme and reaction modalities (Figure 1). For the enzyme modality, GERO uses an amino acid sequence embedding from ESM-2 to summarize global evolutionary and functional signals, and a sequence-derived pocket graph constructed from AlphaFold-predicted structures and P2Rank-predicted ligand-binding pockets to capture local geometric context. For the reaction modality, molecular participants are encoded with Uni-Mol and pooled into a reaction-level representation. These enzyme and reaction representations are projected to a shared latent dimension and fused by a feature-gating module that produces a joint enzyme-reaction representation for three prediction heads: matching score, optimal pH and optimal temperature.

**Figure 1.**
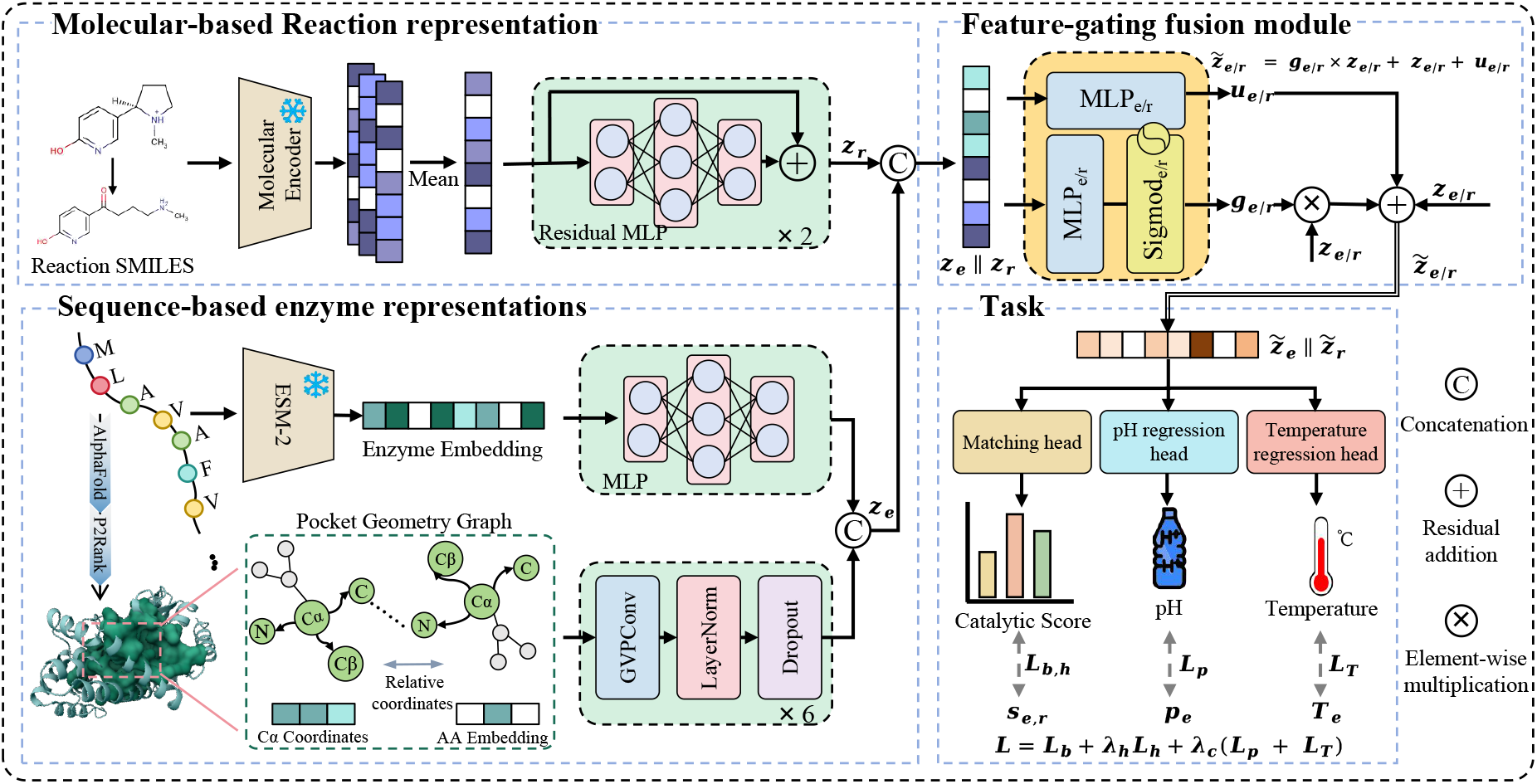
Overview of GERO. Reaction representations are fused with ESM-2 enzyme embeddings and sequence-derived pocket geometry to predict enzyme-reaction matching scores, optimal pH and temperature.

### Molecular-based reaction representation

Reactions are encoded from their molecular participants because the reactants and products directly define the chemical transformation that an enzyme must recognize and catalyze. GERO first encodes each molecular participant, aggregates the participant embeddings into an order-invariant reaction vector and then projects this vector into the same latent space as the enzyme representation for cross-modal matching and optima prediction.

Let 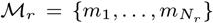 denote the set of available molecular participants in reaction *r*, including reactants and products. Each participant is encoded by a pretrained Uni-Mol molecular encoder. The initial reaction representation **r**_rxn_ is obtained by order-invariant mean pooling:

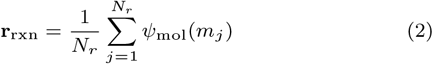

where *ψ*_mol_(·) denotes the molecular encoder. This aggregation provides a compact representation of the chemical participants while avoiding dependence on an arbitrary ordering of molecules in the reaction record.

The pooled reaction embedding is projected into the same latent dimension as the enzyme representation using a residual feed-forward module:

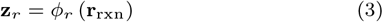

where *ϕ*_*r*_ denotes the reaction projection network. The resulting reaction representation is used in the subsequent feature-gating fusion module.

### Sequence-based enzyme representations

The enzyme representation is designed to capture two related but distinct aspects of retrieval: global catalytic function and local structural context. Global sequence representations capture evolutionary and functional signals that determine broad enzyme-reaction compatibility, whereas local pocket geometry provides structural context that may be relevant to substrate recognition and physicochemical constraints associated with catalytic conditions.

Given an amino acid sequence 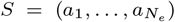, residue-level embeddings are obtained from a pretrained ESM-2 protein language model. The sequence representation is computed by mean pooling over non-special residues:

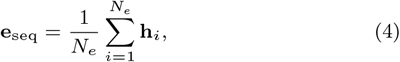

where **h**_*i*_ is the final-layer contextual embedding of residue *i*. This branch provides a global summary of evolutionary and functional information.

To add local structural context without requiring experimentally resolved structures, we applied P2Rank to AlphaFold-predicted enzyme structures to identify putative ligand-binding pockets. The top-ranked predicted pocket was used to define a local pocket graph, providing a structural proxy for the putative catalytic-pocket environment. Let **c** be the predicted pocket center and **x**_*i*_ be the C_*α*_ coordinate of residue *i*. The pocket node set is defined as:

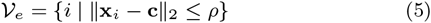

where *ρ* = 15 Å in this study. Edges are defined by a *k*-nearest-neighbor graph over the selected C_*α*_ coordinates:

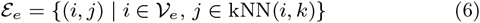

Each node is initialized with residue identity and local orientation vectors derived from backbone atoms and side-chain direction. The resulting pocket graph 𝒢_*e*_ = (𝒱_*e*_, ℰ_*e*_) is encoded by a geometric vector perceptron network (Jing et al., 2020):

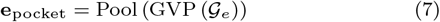

The sequence and pocket embeddings are then concatenated and projected to obtain the final enzyme representation:

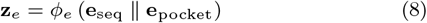

where ∥ denotes concatenation and *ϕ*_*e*_ is a learnable projection network.

### Feature-gating fusion module

After obtaining enzyme and reaction representations, GERO uses feature-gated fusion to build a pair representation for compatibility scoring and condition prediction. Unlike simple concatenation, this module lets features from one modality modulate the other before the matching, pH and temperature heads.

Given **z**_*e*_ and **z**_*r*_, we first form a joint context vector:

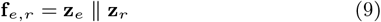

The context vector is used to generate modality-specific gates:

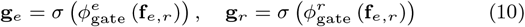

These gates allow the paired reaction to modulate enzyme features and the paired enzyme to modulate reaction features.

To further capture feature-wise compatibility, we compute interaction embeddings from the element-wise product:

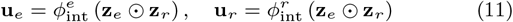

where ⊙ denotes element-wise multiplication. The modulated enzyme and reaction representations are then computed as:

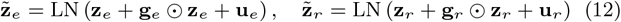

where LN(*·*) denotes layer normalization. The final pair representation is:

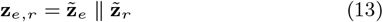

The same joint enzyme-reaction representation is used for all prediction heads:

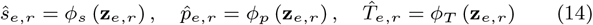

Here, *ŝ* _*e,r*_ is used for candidate ranking, whereas 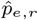 and 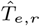 are used to evaluate condition-estimation accuracy under the tolerance criteria.

### Training objective

The training objective is designed to learn a discriminative enzyme-reaction retrieval space while using catalytic-condition annotations where they are available. For a minibatch of *B* positive enzyme-reaction pairs 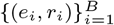, basic in-batch negatives are constructed by randomly permuting the reaction indices, providing a basic contrastive signal for distinguishing observed catalytic associations from random mismatches. The basic matching loss is:

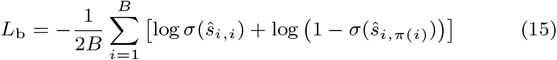

where *π*(·) denotes a random permutation with *π*(*i*) ≠ *i, ŝ*_*i,j*_ is the predicted matching score between enzyme *e*_*i*_ and reaction *r*_*j*_, and *σ*(·) denotes the sigmoid function. This loss encourages observed enzyme-reaction pairs to receive high scores and randomly mismatched pairs from the same minibatch to receive low scores.

To improve discrimination among chemically or functionally similar candidates, we further use in-batch hard negative mining in both retrieval directions. For each enzyme anchor *e*_*i*_, cosine similarities are computed between 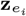 and all reaction embeddings in the minibatch. True positives, identical pairs and candidate pairs with cosine similarity greater than 0.95 are excluded to reduce the risk of selecting false negatives. From the remaining candidates, the top-*K* most similar reactions are selected as the hard negative set 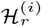, with *K* = 10 in this study. Symmetrically, the same filtering and selection procedure is applied to each reaction anchor *r*_*i*_ to obtain the hard negative enzyme set 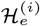. The hard-negative loss is:

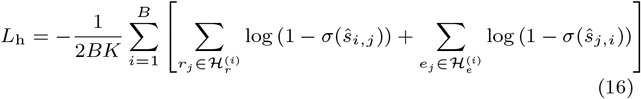

Because pH and temperature annotations are incompletely reported in public databases, the corresponding losses are computed only for positive enzyme-reaction pairs with available condition labels. These labels are treated as optimal conditions, while the paired reaction is used as contextual information to support pH and temperature prediction. Let ℬ_*p*_ and ℬ_*T*_ denote the labeled positive subsets for optimal pH and temperature, respectively. The two condition-prediction losses are:

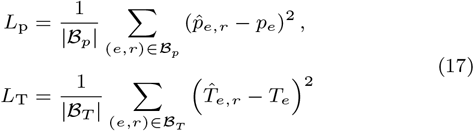

If a minibatch contains no valid annotation for one condition target, the corresponding term is omitted. The total objective combines retrieval and condition supervision:

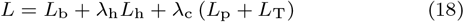

where *λ*_h_ and *λ*_c_ control the relative contributions of hard-negative learning and condition prediction. In practice, condition supervision is introduced after an initial retrieval-alignment stage, allowing the model to first learn a stable enzyme-reaction matching space before optimizing catalytic optima estimates with reaction context.

## Experiments and results

### Experimental setup

#### Dataset construction

We constructed a benchmark by linking reviewed Swiss-Prot (Consortium, 2019) enzyme sequences with Rhea (Bansal et al., 2022) reactions, AlphaFoldDB (Jumper et al., 2021) structures and BRENDA (Placzek et al., 2016) pH and temperature annotations. After preprocessing, the benchmark contained 152,379 enzyme-reaction pairs, 109,254 unique enzymes and 7,572 unique reactions. We then generated two cluster-based evaluation datasets, Enzyme-SIM and Reaction-SIM, by splitting clusters defined by enzyme similarity and reaction similarity, respectively. Further details are provided in Supplementary Section S1.

#### Evaluation settings

We evaluated retrieval using mean reciprocal rank (MRR) and Hit@k, condition regression using mean absolute error (MAE) and root mean squared error (RMSE), and joint condition-aware retrieval using tolerance-restricted Hit@k (Hit@k-TR). Hit@k-TR counts a successful prediction only when the correct candidate appears in the top-*k* list and the predicted pH and temperature fall within predefined tolerances, set here to Δ_*p*_ = 0.5 pH units and Δ_*T*_ = 5°C. Metric definitions are provided in Supplementary Section S2.

Because no existing method jointly performs enzyme-reaction retrieval and catalytic optima prediction, we compared GERO with two-stage baselines that retrieve candidates using SimSearch, ReactZyme (Hua et al., 2024) or CREEP adapted from CARE (Yang et al., 2024), followed by pH and temperature prediction using Seq2pH/Topt (Qiu et al., 2025). We also included GERO*, a decoupled variant that uses GERO for retrieval and Seq2pH/Topt for condition prediction. Baseline implementation details are provided in Supplementary Section S3.

#### Parameter settings

GERO was implemented in PyTorch and trained with the AdamW optimizer. The ESM-2 sequence encoder and Uni-Mol reaction encoder were frozen, and their representations were projected into a 256-dimensional shared latent space. The model was trained for 100 epochs with a batch size of 256 and a base learning rate of 6*×*10^*−*4^. The loss weights were set to *λ*_h_ = 10.0 and *λ*_c_ = 0.75, and the pH and temperature regression losses were activated after epoch 30. Additional training details are provided in Supplementary Section S4.

### GERO improves tolerance-restricted retrieval and prediction performance

We evaluated the E→R and R→E tasks on two distinct datasets to determine whether GERO improves tolerance-restricted enzyme-reaction retrieval performance. Across these settings, GERO achieved the best Hit@k-TR performance (Figure 2). Specifically, in the E→R task on the Enzyme-SIM dataset, GERO achieved Hit@1-TR and Hit@10-TR scores of 24.00% and 35.56%, respectively. These values correspond to absolute gains of 5.33 and 12.89 percentage points over the strongest baselines, SimSearch (18.67%) and CREEP (22.67%). Because Enzyme-SIM is categorized based on enzyme sequence similarity, these results support the model’s ability to identify potential catalytic reactions for novel enzymes. To illustrate this behavior more concretely, we examined the reaction search results for enzyme Q92769 Supplementary Figure S1a. GERO retrieved the correct target reaction and predicted catalytic conditions close to the curated annotations, with maximum deviations of 0.3 pH units and 5.1°C among the annotated examples. Moreover, the retrieved target reaction RHEA:69176 was absent from the training set, further supporting the ability of GERO to generalize to unseen enzyme-reaction pairs. In the R→E task on the same dataset, GERO showed a similar ability to match known reactions to candidate enzymes, achieving a Hit@10-TR of 32.21%.

**Figure 2.**
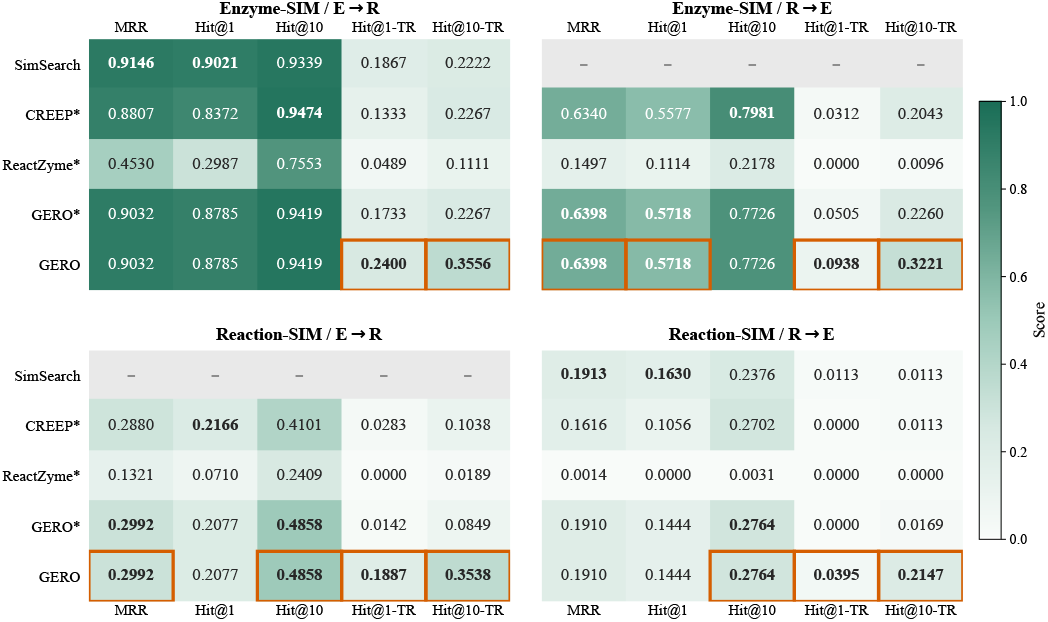
Performance heatmaps on Enzyme-SIM and Reaction-SIM. Darker colors indicate higher scores, and orange boxes highlight the best GERO results.

In the R→E task on the Reaction-SIM dataset, where query reactions are chemically dissimilar from those seen during training, GERO achieved a Hit@10-TR score of 21.47%, indicating its ability to identify suitable catalytic enzymes for new reactions. Notably, the comparative methods all yielded Hit@10-TR scores below 2% in this task. Although the baseline methods retained certain ranking signal, their retrieved candidates are limited by bottlenecks in their condition prediction models. In contrast, GERO, operating as a unified framework, alleviated this bottleneck by integrating condition prediction into the retrieval model. Supplementary Figure S1b presents representative Reaction-SIM examples across both retrieval directions. For example, when retrieving the catalytic enzyme for RHEA:55072, GERO recovered the correct enzyme and predicted its optimal pH and temperature with low deviation from the curated annotations. Additionally, although SimSearch performs well in certain tasks, such similarity-based retrieval methods still face inherent limitations when confronted with unknown reactions (Figure 2).

Finally, to examine whether reaction information provides a richer catalytic context for pH and temperature prediction, we evaluated three enzyme-reaction pairing strategies (random, Top-1 retrieval, random positive). As shown in Figure 3, random reaction pairing did not improve prediction, whereas top-ranked reaction context and diagnostic ground-truth positive context reduced MAE for both pH and temperature compared with the enzyme-only Seq2pH/Topt baseline. The effect was especially clear for temperature prediction on the Enzyme-SIM split, where using the top-ranked reaction context reduced MAE from 7.8202 to 6.8185, corresponding to a 12.81% relative error reduction. These results indicate that informative reaction context contributes to catalytic optima estimation.

**Figure 3.**
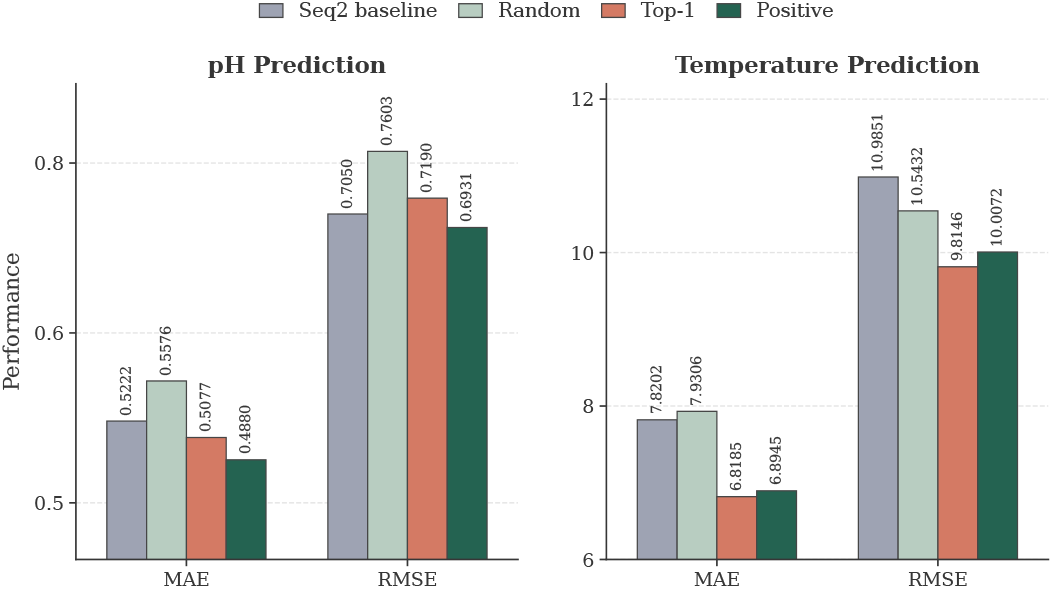
pH and temperature prediction errors under different reaction-context settings, compared with the enzyme-only Seq2pH/Topt baseline.

### Analysis of module contributions and representation quality

To investigate the sources of GERO’s performance, we conducted ablation analyses across benchmark splits and retrieval directions (Figure 4 and Supplementary Figure S2). In the Enzyme-SIM E→R task, removing the pocket-derived structural branch reduced Hit@1-TR from 24.00% to 19.56% and Hit@10-TR from 35.56% to 31.11%, while removing the feature-gated fusion module decreased them to 22.67% and 33.78%, respectively. These results show that pocket-level structural information and feature-gated fusion both contribute to enzyme-reaction matching.

**Figure 4.**
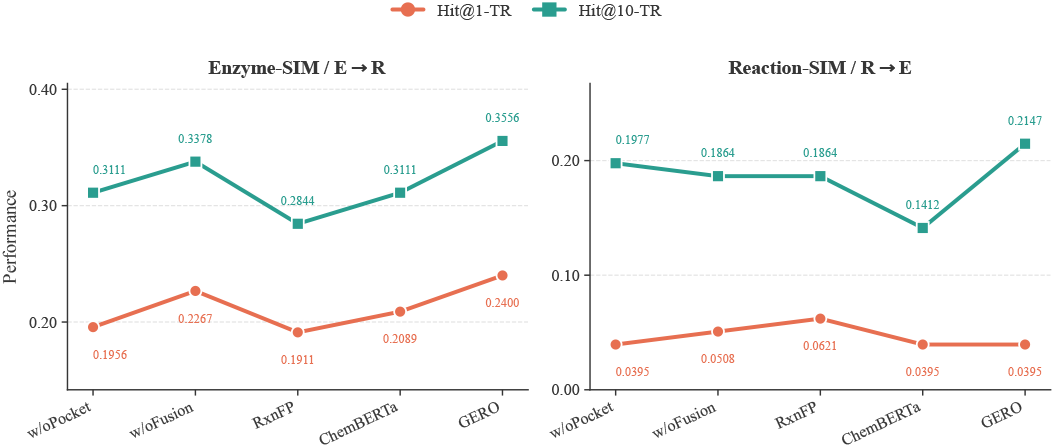
Ablation results for Enzyme-SIM E→R and Reaction-SIM R→E retrieval, comparing pocket, fusion and reaction-encoder variants.

We also assessed the impact of reaction representation by replacing the Uni-Mol reaction encoder with RxnFP (Schwaller et al., 2021) and ChemBERTa (Chithrananda et al., 2020). Both alternatives led to lower performance in the Enzyme-SIM E→R task, with Hit@10-TR decreasing to 28.44% and 31.11%, respectively. In the more challenging Reaction-SIM R→E task, GERO achieved the highest Hit@10-TR of 21.47%, whereas ablated or encoder-replaced variants showed lower top-10 retrieval performance. Together, these results indicate that reaction representation quality, pocket-derived structural cues, and feature-gated fusion jointly support GERO’s retrieval capability.

We further examined the learned representation through score-distribution and t-SNE analyses (Figure 5). The cumulative distributions of alignment scores showed clear separation between positive pairs and random negatives in both the training and test sets: most random negatives scored below 0, whereas positive pairs were concentrated above 0. In t-SNE projections (Van der Maaten and Hinton, 2008) of fused multimodal embeddings on the Enzyme-SIM test set, samples were organized by broad EC class, with EC-2 and EC-3 forming particularly cohesive regions. When samples were grouped by pH using a threshold of 7.4 or by temperature using a threshold of 35°C, locally enriched regions were also observed. These patterns support that the learned representation captures both catalytic matching and condition-related information.

**Figure 5.**
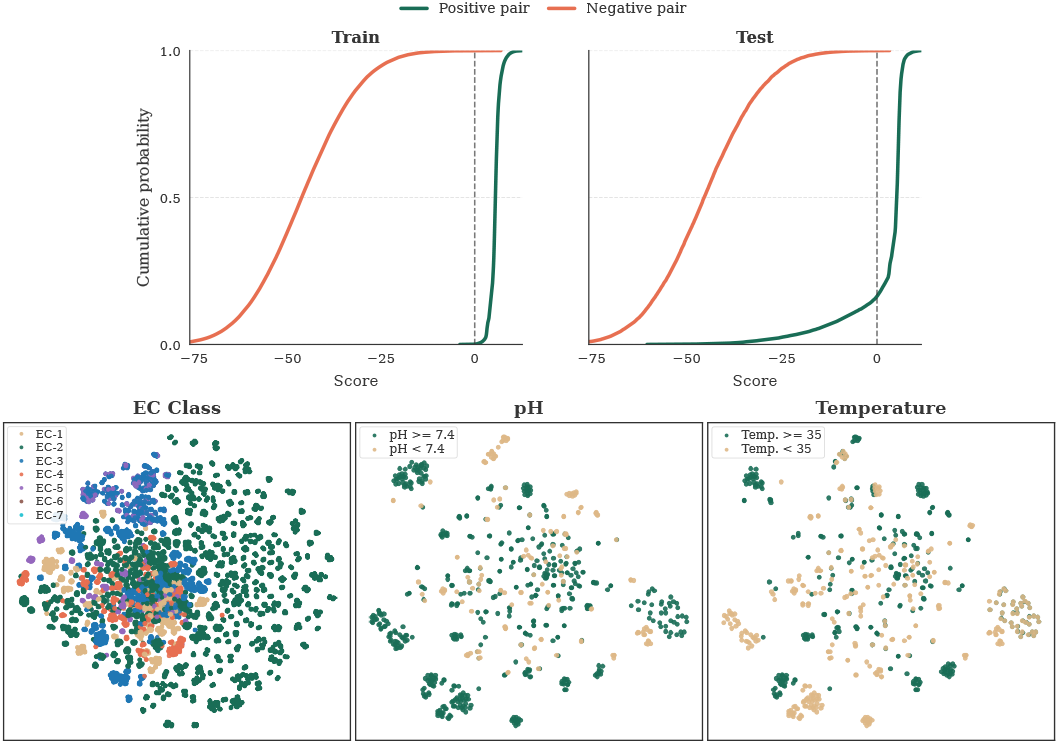
Representation-quality analyses. Top, score distributions for positive and negative enzyme-reaction pairs. Bottom, t-SNE visualization of fused pair representations by EC class, pH group and temperature group.

### Application to iodinin biosynthesis

We next examined whether GERO can support pathway-level enzyme prioritization outside the benchmark retrieval setting. We used the iodinin biosynthesis pathway recently curated in MetaCyc (Caspi et al., 2018) as a case study. Iodinin is a phenazine compound, and its biosynthetic pathway has been characterized through studies in *Lysobacter antibioticus* and *Xenorhabdus szentirmaii* (Zhao et al., 2016; Shi et al., 2019). The pathway starts from phenazine-1,6-dicarboxylate and proceeds through four reactions catalyzed by xpzG, xpzH and phzNO1.

Because these pathway enzymes and reactions were absent from the training set, the case study evaluates whether GERO can prioritize enzymes within a pathway-level candidate pool. We used the model trained on the Reaction-SIM split and constructed a retrieval library of 190 enzymes from relevant UniRef50 clusters with available AlphaFold structures. During inference, reaction inputs were restricted to the primary reactants and products, excluding coenzymes.

GERO placed the known pathway enzymes within the candidate rankings for all four reaction steps (Figure 6). Its prioritization was strongest for the first two reactions, which involve larger scaffold changes; for the second step, where the comparison retrieval methods ranked the target enzyme outside the top candidates, GERO ranked the target enzyme at position 6. In contrast, for the later N-oxidation steps, ReactZyme or CREEP sometimes ranked the target enzymes higher than GERO, suggesting that the current participant-aggregation reaction representation remains less sensitive to transformations dominated by local charge transfer. Because these pathway enzymes do not have standardized optimal pH or temperature annotations in the databases used here, we compared the predicted conditions with assay conditions reported in the literature rather than treating them as ground-truth optima. The xpzG/xpzH assays were performed at pH 8.5, whereas phzNO1 assays were performed at pH 7.5, both at approximately 30°C (Zhao et al., 2016; Shi et al., 2019). The GERO estimates, although showing some pH deviation, remained within practically acceptable tolerance ranges around the reported assay conditions.

**Figure 6.**
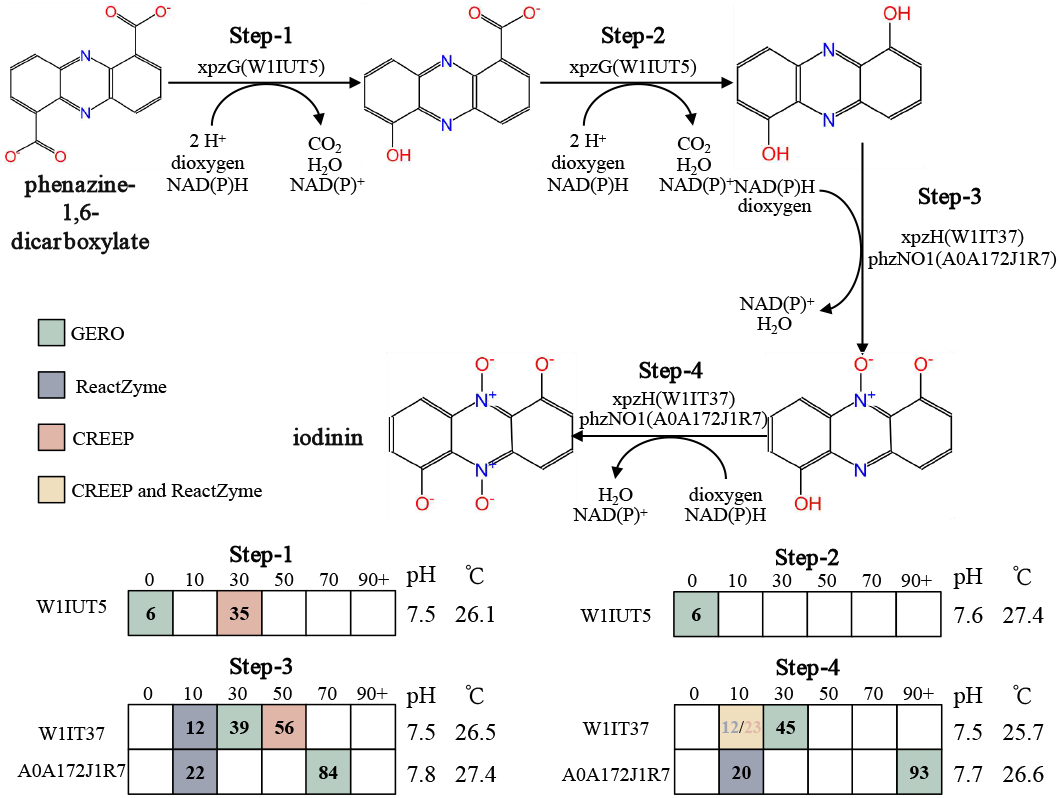
Application to iodinin biosynthesis. The pathway consists of four reaction steps catalyzed by three enzymes. The lower panels pH and temperature values are GERO estimates.

## Conclusion and discussion

We presented GERO, a multimodal fusion framework for condition-aware enzyme-reaction retrieval and catalytic optima estimation with reaction context. By integrating global enzyme sequence semantics, predicted pocket geometry, and molecular reaction representations through feature-gated cross-modal fusion, GERO learns a shared enzyme-reaction representation to jointly rank candidate pairs and estimate likely optimal pH and temperature conditions.

Across both Enzyme-SIM and Reaction-SIM benchmark splits, GERO improved Hit@k-TR over two-stage retrieval-then-prediction baselines. However, several potential limitations remain. First, public pH and temperature annotations are incomplete and are often reported at the enzyme, organism, or EC-associated record level rather than as fully observed enzyme-reaction-specific activity profiles, which may limit the specificity and accuracy of condition prediction for enzyme-reaction pairs. Second, the current reaction representation aggregates molecular participants and does not explicitly model atom mapping, stoichiometry, or reaction-center electron flow, which may reduce sensitivity to transformations driven by subtle local chemical changes. Third, the structural branch relies on P2Rank-predicted binding pockets from AlphaFold-predicted structures rather than experimentally confirmed catalytic sites, which may introduce uncertainty into pocket-level features and limit their interpretation as direct catalytic-site evidence.

Future work could extend condition-aware retrieval beyond ranking within a predefined candidate space toward condition-guided enzyme candidate generation or engineering, where desired reactions and assay conditions guide the proposal of new or modified enzyme candidates. Methodologically, future models could incorporate atom-mapped reaction centers, reaction-difference graphs, docking or complex-structure constraints, and uncertainty estimates. Experimental validation will ultimately be needed to determine whether condition-aware recommendations improve the efficiency of biocatalyst discovery workflows.

## Data availability

Reproducible data and code are available at https://github.com/ykxhs/GERO.

