## Supplementary materials for "Joint enzyme-reaction retrieval and catalytic optima prediction via multimodal fusion"

#### S1. Dataset construction and benchmark splits

**Dataset construction:** To construct a high-quality dataset of enzyme-reaction pairs alongside optimal pH and temperature, we integrated four bioinformatics databases: UniProt, Rhea, BRENDA, and AlphaFold DB. First, we retrieved all non-fragment enzyme entries with Rhea annotations from the UniProt (Swiss-Prot, release 2025.10.26) database, ensuring that the selected enzymes possessed explicit Rhea reaction annotations and were complete, manually reviewed sequences. To accommodate the maximum token length of the ESM-2 pretrained model, we filtered out enzyme entries with sequence lengths exceeding 1022. The 3D structural data for all enzymes were mapped and retrieved from the AlphaFold DB. Second, based on the retrieved Rhea identifiers, we obtained the corresponding reaction SMILES from Rhea (release 139), uniformly adopting the left-to-right direction as the standardized representation of the reactions. All reactions were subjected to impurity removal, metal disconnection, and charge neutralization. By eliminating disparities in molecular states caused by varying experimental environments, we converted them into unique and canonical representations.

Furthermore, because BRENDA (release 2025.1) organizes data by EC numbers, we iterated through all EC classes to search for entities with documented optimal pH and temperature values. For data provided as ranges, we calculated the mean. For entities with multiple data records, we discarded entries where the pH difference exceeded 1 or the temperature difference exceeded 10°C. Ultimately, the average of the remaining data was assigned as the optimal pH and temperature for the enzyme. The final constructed dataset comprises 152,379 enzyme-reaction pairs, encompassing 109,254 unique enzyme sequences and 7,572 unique chemical reactions. Among these, 2,851 enzymes possess optimal pH annotations, and 2,435 enzymes possess optimal temperature annotations. Supplementary Table S1 reports the marginal availability of pH and temperature annotations. **Benchmark splits:** We used two cluster-based splits to evaluate out-of-distribution generalization. For the enzyme split, MMseqs2 clustered enzyme sequences at 50% sequence identity. For the reaction split, reactions were represented by averaged ECFP6 fingerprints of their molecular participants and clustered with the Butina algorithm using a distance threshold of 0.5. Whole clusters were assigned to training or test sets at an approximate 9:1 ratio, ensuring that close homologs or chemically similar reactions did not appear in both sets.

For enzyme-to-reaction retrieval, each enzyme was ranked against all unique candidate reactions in the benchmark. For reaction-to-enzyme retrieval, each reaction was ranked against all unique candidate enzymes. In the iodinin biosynthesis case study, the enzyme candidate pool was limited to 190 enzymes from UniRef50 clusters associated with the target pathway enzymes and with available AlphaFold structures. Reaction inputs in this case study included only primary reactants and products, excluding coenzymes so that retrieval focused on the core chemical transformation.

#### S2. Evaluation metrics

Retrieval performance was evaluated in both enzyme-to-reaction and reaction-to-enzyme directions. For a query $q$, candidates from the opposite modality were ranked in descending order according to the predicted matching score $s(q, c)$. Let $\mathcal{G}(q)$ denote the set of ground-truth candidates for query $q$. The rank of the first correct candidate was defined as

$$\mathrm{rank}(q) = \min_{c \in \mathcal{G}(q)} \mathrm{rank}_q(c), \tag{1}$$

where $\mathrm{rank}_q(c)$ is the rank position of candidate $c$ for query $q$. Hit@k was computed as

$$\mathrm{Hit@}k = \frac{1}{|Q|} \sum_{q \in Q} \mathbf{1}\left(\mathrm{rank}(q) \le k\right), \tag{2}$$

and mean reciprocal rank (MRR) was computed as

$$\mathrm{MRR} = \frac{1}{|Q|} \sum_{q \in Q} \frac{1}{\mathrm{rank}(q)}. \tag{3}$$

The tolerance-restricted hit rate, Hit@k-TR, jointly evaluates candidate retrieval and catalytic-condition prediction. Let $P \subseteq Q$ denote the subset of queries for which at least one ground-truth enzyme-reaction pair has both pH and temperature annotations, and let $\mathcal{G}_{pt}(q) \subseteq \mathcal{G}(q)$ denote the annotated ground-truth candidates for query $q$. For a retrieved ground-truth candidate $c$, the tolerance condition was defined as

$$\mathcal{T}(q, c) = \max\left(\frac{|\hat{y}_p(q, c) - y_p(q, c)|}{\Delta_p}, \frac{|\hat{y}_T(q, c) - y_T(q, c)|}{\Delta_T}\right) \le 1, \tag{4}$$

where $\hat{y}_p$ and $\hat{y}_T$ denote predicted optimal pH and temperature, $y_p$ and $y_T$ denote annotated values, and $\Delta_p$ and $\Delta_T$ denote the predefined tolerance thresholds. The set of tolerance-satisfying retrieved ground-truth candidates was

$$\mathcal{H}_k^{\mathrm{TR}}(q) = \left\{c \in \mathrm{Top}_k(q) \cap \mathcal{G}_{pt}(q) \mid \mathcal{T}(q, c)\right\}. \tag{5}$$

Hit@k-TR was then computed as

$$\text{Hit@}k\text{-TR} = \frac{1}{|P|} \sum_{q \in P} \mathbf{1} \left[ \mathcal{H}_k^{\text{TR}}(q) \neq \emptyset \right]. \tag{6}$$

In all experiments, $\Delta_p$ was set to 0.5 pH units and $\Delta_T$ was set to 5°C.

Continuous pH and temperature prediction was evaluated using mean absolute error (MAE) and root mean squared error (RMSE) on the corresponding labeled subsets. For target $a \in \{p, T\}$ and labeled test subset $\mathcal{D}_a$, these metrics were defined as

$$\text{MAE}_a = \frac{1}{|\mathcal{D}_a|} \sum_{i \in \mathcal{D}_a} \left| y_a^{(i)} - \hat{y}_a^{(i)} \right|, \tag{7}$$

$$\text{RMSE}_a = \sqrt{\frac{1}{|\mathcal{D}_a|} \sum_{i \in \mathcal{D}_a} \left( y_a^{(i)} - \hat{y}_a^{(i)} \right)^2}. \tag{8}$$

#### S3. Baseline implementation details

Because no existing method jointly performs bidirectional enzyme-reaction retrieval and catalytic optima prediction, all comparison methods were implemented as two-stage baselines. Each retrieval baseline first ranked candidate enzymes or reactions, and Seq2pH/Topt was then used to provide enzyme-level optimal pH and temperature predictions for the retrieved candidates. This design mirrors the common decoupled workflow and provides a direct comparison against the training strategy used by GERO, which incorporates reaction context.

SimSearch was used as a non-parametric similarity-search baseline. For enzyme queries, MMseqs2 was used to compute amino acid sequence identity against the training set. Reactions associated with the matched training enzymes were assigned to the query, and the maximum sequence-identity score was retained when multiple homologous enzymes mapped to the same reaction. For reaction queries, each reaction was represented by averaged ECFP6 fingerprints of its molecular participants, and Tanimoto similarity was computed against training-set reactions. Because the benchmark splits were cluster-based, this baseline was evaluated only in settings where the similarity search was meaningful under the corresponding split: enzyme-to-reaction retrieval for the Enzyme-SIM split and reaction-to-enzyme retrieval for the Reaction-SIM split.

ReactZyme was reproduced from its open-source implementation (`https://github.com/WillHua127/ReactZyme`) using the Uni-Mol+ESM+NoGNN architecture. Because the released repository did not provide a complete negative-sampling script, negative samples were generated following the original paper, with 1,000 negative samples constructed for the enzyme side and 1,000 negative samples constructed for the reaction side during training. To improve data-loading efficiency on the present benchmark, sequence and reaction SMILES storage keys were replaced with unique identifiers. The model was retrained on the benchmark using a single NVIDIA RTX 4090 GPU.

CREEP was reproduced from the CARE benchmark implementation (`https://github.com/jsunn-y/CARE`). The original tri-modal enzyme-reaction-text alignment architecture was adapted by removing the text encoder and using the dot product between enzyme and reaction embeddings as the retrieval score. To keep the protein encoder consistent across baselines, the original ProtT5 encoder was replaced with ESM-2 (650M). The model was trained with the BM-NCE loss and otherwise retained the default hyperparameter settings from the released implementation. Retraining was performed using four NVIDIA RTX 4090 GPUs.

Seq2pH/Topt was used as the enzyme-level catalytic optima predictor in the two-stage baselines. The model predicts optimal pH and temperature from amino acid sequence inputs using protein language-model features and residual prediction heads. For the present comparison, Seq2pH/Topt was reproduced from the open-source implementation (`https://github.com/SizheQiu/Seq2Topt`) and retrained on the present benchmark using a single NVIDIA RTX 4090 GPU. To isolate the contribution of joint condition-aware training, a decoupled GERO variant, denoted GERO*, used GERO for retrieval but replaced its pH and temperature estimates with Seq2pH/Topt outputs.

#### S4. Training and implementation settings

GERO was implemented in PyTorch and trained with AdamW. ESM-2 (650M) and Uni-Mol were used as the protein and molecular encoders, respectively. Their parameters were frozen during training to preserve pretrained representations and reduce computational cost. AlphaFoldDB version 6 provided predicted structures, and P2Rank version 2.5.1 was used for pocket identification. Only the downstream geometry-aware graph network, projection layers, feature-gating module, and task-specific heads were updated.

For each enzyme, the ESM-2 sequence embedding was obtained by mean pooling the final-layer residue embeddings, excluding special tokens and padding positions. The original ESM embedding dimension was 1280. For the pocket branch, the top-ranked P2Rank pocket was used as the pocket center. Residues with $C_\alpha$ atoms within 15 Å of this center were selected as graph nodes. Each node included an amino acid identity feature and local orientation vectors from $C_\alpha$ to the backbone N atom, backbone C atom, and $C_\beta$ atom. For glycine, the $C_\beta$ direction was approximated using the normalized cross product of backbone direction vectors. Directed edges were built with a $k$-nearest-neighbor graph using $k = 20$, based on $C_\alpha$ distances. Each edge carried the relative displacement vector between the connected residues.

The pocket graph was encoded with a Geometric Vector Perceptron graph neural network. Amino acid identities were mapped to learnable scalar embeddings, and vector features were initialized from the local orientation vectors. A preliminary GVP layer first aligned scalar and vector dimensions, followed by a six-layer GVP graph convolutional network with residual connections and dropout. The final scalar node states were normalized, passed through a readout multilayer perceptron, and mean pooled to produce the pocket embedding.

This pocket embedding was concatenated with the projected ESM sequence embedding and then linearly projected to form the enzyme representation.

For each reaction, all available reactants and products were encoded with Uni-Mol and combined by order-invariant mean pooling. The original Uni-Mol reaction embedding dimension was 512. The pooled reaction embedding was projected into the fusion dimension and processed by two pre-norm residual feed-forward blocks. Each block contained layer normalization, a SiLU-activated feed-forward network, and a residual connection. The final reaction representation was layer-normalized before fusion. Both enzyme and reaction representations were mapped to a shared 256-dimensional latent space.

The feature-gating module generated modality-specific gates from the concatenated enzyme and reaction representations. It also computed interaction features using the element-wise product of the two representations. For each modality, the gated residual update combined the original representation, the gate-modulated representation, and the interaction feature, followed by layer normalization. The updated enzyme and reaction representations were then concatenated as the joint pair representation. Three independent two-layer prediction heads with ReLU activation were used to predict the matching logit, optimal pH, and optimal temperature.

Hard-negative mining was performed within each minibatch in both retrieval directions. For each enzyme anchor $e_i$, cosine similarities were computed between $\mathbf{z}_{e_i}$ and all reaction embeddings in the minibatch. True positives, identical pairs, and candidate pairs with cosine similarity greater than 0.95 were excluded to reduce the risk of selecting false negatives. From the remaining candidates, the top-$K$ most similar reactions were selected as $\mathcal{H}_r^{(i)}$, with $K = 10$. The same filtering and selection procedure was applied to each reaction anchor $r_i$ to obtain the hard negative enzyme set $\mathcal{H}_e^{(i)}$.

The model was trained for 100 epochs with a batch size of 256 on a single NVIDIA RTX 4090 GPU. The base learning rate was $6 \times 10^{-4}$, and weight decay was $1 \times 10^{-5}$. Training used a 20-epoch linear warm-up from a factor of 0.01, followed by cosine annealing to a minimum learning rate of $1 \times 10^{-6}$. Gradients were clipped at a maximum norm of 1.0. The loss weights were $\lambda_{\text{hard}} = 10.0$ and $\lambda_{\text{cond}} = 0.75$. Condition-regression loss was introduced at epoch 30, allowing the model to first learn enzyme-reaction alignment before optimizing pH and temperature prediction.

### Supplementary figures

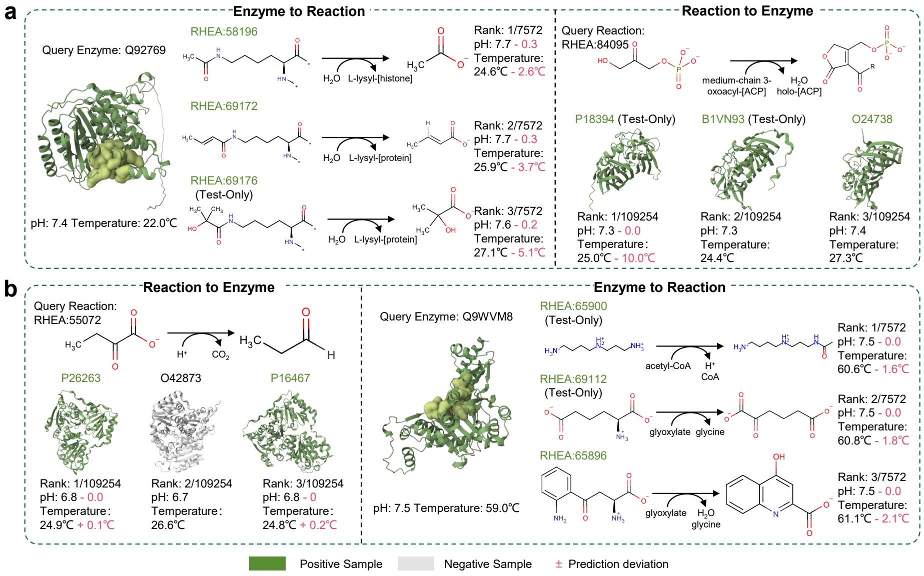


Figure S1: Representative benchmark retrieval examples. a, Top-ranked candidate reactions for query enzyme Q92769 from the Enzyme-SIM test set and top-ranked candidate enzymes for query reaction RHEA:84095. b, Top-ranked candidate enzymes for query reaction RHEA:55072 from the Reaction-SIM test set and top-ranked candidate reactions for query enzyme Q9WVM8. Predicted pH and temperature values are shown for each retrieved candidate, with deviations from curated annotations indicated when available.

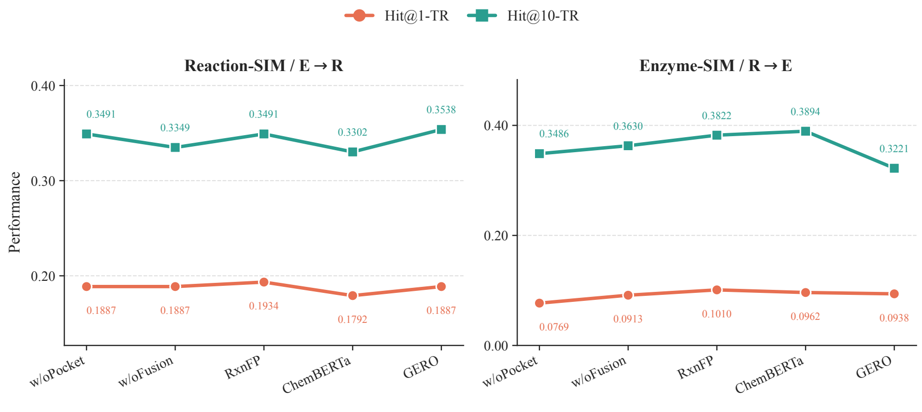


Figure S2: Ablation results for Reaction-SIM E→R and Enzyme-SIM R→E retrieval, comparing pocket, fusion and reaction-encoder variants.

#### Supplementary tables

Table S1: Detailed sample counts for the Enzyme-SIM and Reaction-SIM benchmark splits.

| | Enzyme-SIM | | Reaction-SIM | |
|---|---|---|---|---|
| **Category** | **Train** | **Test** | **Train** | **Test** |
| Enzyme-reaction pairs | 136,658 | 15,721 | 142,204 | 10,175 |
| Unique enzymes | 98,344 | 10,910 | 102,908 | 7,803 |
| Unique reactions | 7,124 | 1,768 | 6,928 | 644 |
| Unique enzymes with pH annotations | 2,561 | 290 | 2,669 | 304 |
| Unique enzymes with temperature annotations | 2,171 | 263 | 2,288 | 252 |

Table S2: Number of eligible test queries used as denominators for Hit@k-TR. Eligible queries are those with at least one ground-truth enzyme-reaction pair containing both pH and temperature annotations.

| **Dataset** | **E→R** | **R→E** |
|---|---|---|
| Enzyme-SIM | 225 | 416 |
| Reaction-SIM | 212 | 177 |
